# Proteome and microbiota analysis reveals alterations of liver-gut axis under different stocking density of Peking ducks

**DOI:** 10.1101/335570

**Authors:** Yuqin Wu, Jianhui Li, Xin Qin, Shiqiang Sun, Zhibin Xiao, Xiaoyu Dong, Muhammad Suhaib Shahid, Dafei Yin, Zhao Lei, Yuming Guo, Jianmin Yuan

## Abstract

The aim of this study was to determine the impact of stocking density on the liver proteome and cecal microbiota of Peking ducks. A total of 1,200 ducks with 21-day old were randomly allotted into 5 stocking density groups of 5, 6, 7, 8 and 9 ducks/m^2^, with 6 replicates for each group. At 40 days of age, duck serum and pectorals were collected for biochemical tests; liver and cecal contents of ducks were gathered for proteome and microbiota analysis, respectively. Serum MDA increased while pectorals T-AOC reduced linearly with enhancing stocking density. Duck lipid metabolism was altered under different stocking density as well. Serum LDL-C increased linearly with increasing stocking density. Proteome analysis revealed fatty acid biosynthesis proteins such as acyl-CoA synthetase family member 2 and fatty acid oxidation related proteins including acyl-CoA dehydrogenase long chain and acyl-coenzyme A oxidase were enriched in high stocking density group. Additionally, high stocking density increased oxidative response related proteins such as DDRGK domain containing 1 while diminished anti-oxidant capacity related proteins including regucalcin and catalase. 16S rDNA analysis revealed that higher stocking density was accompanied with decreased microbial diversity, as well as depletion of anti-inflammatory bacterial taxa, including *Bacteroidales, Butyricimonas* and *Alistipe*. In addition, decreased bile acid metabolism-associated bacteria such as *Ruminococcaceae, Clostridiales* and *Desulfovibrionaceae* were found in the high-density group. Both proteome and 16S rDNA results showed inflammation and chronic liver disease trend in the high-density group, which suggests the involvement of the liver-gut axis in oxidative stress.

## INTRODUCTION

Space is one of the most compromised features in commercial housing systems, limited often in the interests of efficiency and profitability. Increasing stocking density can yield higher profits per kilogram of chicken. However, reduced space limits the ability to rest [1] and has a negative influence on performance [2,3,4,5,6], meat yield [2-4], immune status [7] and gut morphology [8]. In addition, high stocking density is associated with chronic oxidative stress [2].

The liver plays an important role in energy metabolism and it is the major site of triglyceride (TG) metabolism which is involved in TG digestion, absorption, synthesis, decomposition and transport. It has been demonstrated that high stocking density has been shown to have a deleterious effect on liver function [9] with increased activities with aspartate aminotransferase and alanine aminotransferase [10]. The gut microbiome is also highly connected to animal energy metabolism and health [11], and it has been frequently suggested that gut microbiota plays a critical role in chronic liver disorders through liver-gut axis [12]. Moreover, high stocking density is associated with adverse effects on the chicken intestinal commensal bacteria [4].

The emergence of novel proteomic techniques in recent years has greatly aided in the understanding of biological mechanisms. Tandem mass tag (TMT) [13] and isobaric tags for relative and absolute quantitation (iTRAQ) [14] methods have been widely used for analyzing the hepatic proteome. The TMT method can also be used to characterize liver proteome-wide changes in response to oxidative stress [15]. Further, due to progress in high-throughput next-generation sequencing, 16S rDNA analysis can be used to infer the structure and function of gut microbiota.

To characterize the potential mechanism of oxidative stress under high stocking density, TMT-labeled quantitative proteomics combined with 16S rDNA analysis was used to identify changes in the protein profiles and microbiota of Peking ducks under low and high raising density.

## MATERIALS AND METHODS

### Ethics approval

All of our experiments were approved by the Institutional Animal Care and Use Committee of the China Agricultural University (Beijing, China).

### Facilities and Experimental Animals

A total of 1200 mixed-sex 21-d-old white Peking ducks were randomly assigned to 5 stocking density treatments and 6 replicates. Each replicate corresponds one pen with 40 ducks (20 males, and 20 females). The raising area in different treatments were 8.20 m^2^ (2.50 × 3.28 m), 6.88 m^2^ (2.50 × 2.75 m), 5.93 m^2^ (2.50 × 2.37 m), 5.20 m^2^ (2.50 × 2.08 m) and 4.65 m^2^ (2.5 × 1.86 m) and the corresponding stocking density was 5 (low stocking density represent as L group), 6, 7, 8, and 9 (high stocking density represent as H group) ducks/m^2^, respectively. All ducks were raised in a plastic wire-floor pen and provided water and feed *ad libitum* from 21 to 40 d of age. In the house, lighting, temperature and ventilation programs followed commercial practices. During the experimental period, all ducks were raised with diets based on the NRC (1994) feeding standard.

### Growth Performance and Carcass Traits

Initial body weight (BW), final BW and feed intake (FI) were recorded. The weight gain and feed/gain ratio were calculated.

### Sample Collection

On day 40, one male duck per pen was randomly selected. Therefore, a total of 6 male ducks per treatment were collected. Each duck was sacrificed by stunning after blood collection. The liver samples of ducks from L and H density treatments were collected, flushed with cold PBS, frozen using liquid nitrogen, and stored at −80 °C for and proteomic analysis. Cecum contents of ducks from L and H density treatments were collected, transferred into Eppendorf tubes, and immediately frozen in liquid nitrogen and stored at −80°C for microbiota analyses.

Abdominal fat and left pectorals of each duck were removed manually from the carcass and weighed. For each duck, a piece of pectorals fixed position was separated and put on the ice bag immediately, then stored at −80 °C for biochemical analysis.

### Serum and Pectorals Biochemical Parameters

Serum levels of malondialdehyde (**MDA**, cat#A003-1), TG (cat#A110-2), total cholesterol (**TC**, cat#A111-2), high-density lipoprotein cholesterol **(HDL-C**, cat#A112-2), low-density lipoprotein cholesterol (**LDL-C**, cat#A113-2), very low density lipoprotein (**VLDL**, cat#H249), total antioxidant capacity (**T-AOC**, cat#A015-1) and the activities of lactate dehydrogenase (**LDH**, cat#A020-1), creatine kinase (**CK**, cat#A032), glutamic-pyruvic transaminase (**GPT**, cat#C009-1), lipase (cat#A067), lipoprotein lipase (**LPL**, cat#A067) along with MDA, T-AOC, LPL and protein concentration (cat#A045-2) of pectorals were determined by using commercial analytical kits according to the manufacturer’s recommendations (Jian Cheng Bioengineering Institute, Nanjing, China).

### Statistical analysis

One-way analysis of variance (ANOVA) models were fitted to assess the relationships between the stocking density groups and serum and pectoral redox and lipid metabolism indices, using SPSS (v.20.0, SPSS Institute, Chicago, IL). Means were compared using Duncan’s multiple comparison procedure of SPSS software when density treatment was significant (*P* < 0.05), and curve estimation was used to assess the linear and quadratic effects of increasing stocking density on final body weight, body weight gain, feed gain ratio, and pectorals percentage. *P* < 0.05 was used for statistical significant and results were considered significant trend at *P* < 0.1.

### Proteome analysis

#### Protein extraction

Liver samples (∼100mg each) were ground to a powder under liquid nitrogen and then transferred into a centrifuge tube. After that, four times volume of lysis buffer containing 8 M urea (Sigma) and 1% Protease Inhibitor Cocktail (Calbiochem) was added, followed by sonication three times on ice using a high intensity ultrasonic processor (Scientz). And then centrifuged for 10 min at 4 °C and 12,000g. Supernatants containing soluble proteins were collected and the protein concentration was quantified with BCA kit (Beyotime Biotechnology) according to the manufacturer’s instructions.

#### Trypsin digestion

For trypsin digestion, the protein solution was reduced with 5 mM dithiothreitol (Sigma) for 30 min at 56 °C and alkylated with 11 mM iodoacetamide (Sigma) for 15 min at room temperature under dark conditions. The protein sample was diluted with 100 mM triethylammonium bicarbonate (TEAB, Sigma) to make urea concentration less than 2 M. Finally, the sample was digested with trypsin (Promega) overnight with a 1:50 trypsin-to-protein ratio and then further digested for 4 h with 1:100 w: w trypsin-to-protein ratio (37℃).

#### TMT Labeling

After trypsin digestion, the peptides were desalted in a Strata X C18 SPE column (Phenomenex) and vacuum dried. The peptide was dissolved in 1 M TEAB and processed according to the manufacturer’s protocol for TMT kit. Briefly, one unit of TMT reagent (defined as the amount of reagent requirement for labeling 100 μg of protein) was thawed and dissolved in acetonitrile (Fisher Chemical). The peptide mixtures were then incubated for 2 h at room temperature, pooled, desalted and dried by vacuum centrifugation.

#### HPLC fractionation

Peptides were fractionated by high pH reverse-phase HPLC with an Agilent 300 Extend C18 column (5 μm particles, 4.6 mm ID, 250 mm length). Briefly, peptides were firstly separated by 8% to 32% acetonitrile (pH 9.0) over 60 minutes into 60 fractions. Subsequently, the peptides were combined into 18 fractions and dried by vacuum centrifugation.

#### Mass Spectrometry and TMT Data Analysis

Peptides were dissolved in 0.1% formic acid (Fluka), and separated by EASY-nLC 1000 UPLC system. Solvent A is an aqueous solution containing 0.1% formic acid and 2% acetonitrile; solvent B is an aqueous solution containing 0.1% formic acid and 90% acetonitrile. Liquid-phase gradient using a linearly increasing gradient of 5% to 24% solvent B for 38 min, 24% to 35% solvent B for 14 min, then climbing to 80% in 4 min, and then maintaining at 80% for the last 4 min, all at a constant flow rate of 800 nl/min.

The peptides results were subjected to nano electrospray ionization (NSI) source followed by tandem mass spectrometry (MS/MS) in Q Exactive PlusTM (Thermo Fisher Scientific) coupled to the UPLC. The electrospray voltage applied was 2.0 kV. For MS scans, the m/z scan range was 350 to 1800. Intact peptides were detected in the orbitrap at a resolution of 70,000. Peptides were then selected for MS/MS in the orbitrap at a resolution of 17,500. Fixed first mass was set as 100 m/z. A data-dependent procedure that alternated between one MS scan followed by 20 MS/MS scans. In order to improve the effective utilization of mass spectrometry, the automatic gain control (AGC) is set to 5E4, the signal threshold is set to 10000 ions/s, the maximum injection time is set to 200 ms, and the dynamic exclusion time of the tandem mass scan is set to 30 seconds to avoid the repeat the scan of parent ion.

#### Database Search

The resulting MS/MS data were processed using Maxquant with an integrated Andromeda search engine (v.1.5.2.8). Tandem mass spectra were searched against the uniprot *Anas platyrhynchos* database concatenated with reverse decoy database. Trypsin/P was specified as cleavage enzyme allowing up to 2 missing cleavages. The mass tolerance for precursor ions was set as to 10 ppm in First search and 5 ppm in Main search, and the mass tolerance was set as 0.02 Da for fragment ions. TMT 10-plex was selected for protein quantification. False discovery rate (FDR) was adjusted to < 1% for protein identification.

#### TMT quantification

For TMT quantification, the ratios of the TMT reporter ion intensities in MS/MS spectra (m/z 126–131) from raw data sets were used to calculate fold changes between samples. For each sample, the quantification was mean-normalized at peptide level to center the distribution of quantitative values. Protein quantitation were then calculated as the median ratio of corresponding unique or razor peptides for a given protein. Two-sample, two-sided T-tests were used to compare expression of proteins. A significance level of *P* < 0.05 was used for statistical testing and results were considered significant trend at *P* < 0.1.

#### Gene Ontology (GO) annotation

GO annotation proteome was derived from the UniProt-GOA database (www. http://www.ebi.ac.uk/GOA/). Firstly, Converting identified protein ID to UniProt ID and then mapping to GO IDs by protein ID. If some identified proteins were not annotated by UniProt-GOA database, the InterProScan soft would be used to annotated protein’s GO functional based on protein sequence alignment method. Then proteins were classified by Gene Ontology annotation based on three categories: biological process, cellular component and molecular function.

### 16S rDNA sequencing

DNA was extracted from 180-220 mg of the cecal samples using a QIAampTM Fast DNA Stool Mini Kit (Qiagene, No. 51604) according to the manufacturer’s instructions. Total DNA was quantified using a Thermo NanoDrop 2000 UV microscope spectrophotometer and 1% agarose gel electrophoresis. 16S rDNA high-throughput sequencing was performed by Realbio Genomics Institute (Shanghai, China) using the Illumina Hiseq PE250 platform. The V3-V4 region of the 16S rDNA gene was amplified using the universal primers, 341F (CCTACGGGRSGCAGCAG) and 806R (GGACTACVVGGGTATCTAATC). The raw pair-end reads were merged and quality-filtered to remove tags with lengths < 220 nt, an average quality score of <20, and tags containing >3 ambiguous bases using PANDAseq (v2.9) [16]. Singletons and chimeras were removed, and the resulting quality-filtered sequences were clustered into 97% operational taxonomic units (OTUs) using USEARCH (v7.0.1090) in QIIME software. The Ribosomal Database Project (RDP) algorithm trained on the Greengenes database was used to classify each OTU (http://greengenes.lbl.gov). The open source software package QIIME (http://qiime.org) was used to measure alpha diversity (including the chao1, observed species and PD whole tree indices).

#### 16S rDNA analysis

For 16S rDNA, the Wilcoxon rank sum test to evaluate changes in alpha diversity between H and L groups. Venn diagrams were constructed in R v3.1.0 using the Venn Diagram package. Principal component analysis (**PCA**) was used to assess the relationships between samples based on composition of the microbiota [17]. Linear discriminant analysis effect size (**LEfSe**), a metagenomic biomarker discovery approach, was used to identify differentially abundant taxa between H and L groups by using a Kruskal-Wallis rank sum test *P*-value threshold of 0.05 and a log-transformed linear discriminant analysis (**LDA**) score threshold of 2.

## RESULTS

### Duck performance and carcass traits

In our study, both the final body weight and the body weight gain over the study period decreased linearly with the increase of stocking density (Fig 1a, 1b). The feed/gain ratio increased linearly with increasing density (Fig 1c). Pectorals percentage was affected by stocking density as well, decreasing linearly with increasing stocking density (Fig 1d).

**Fig 1.**
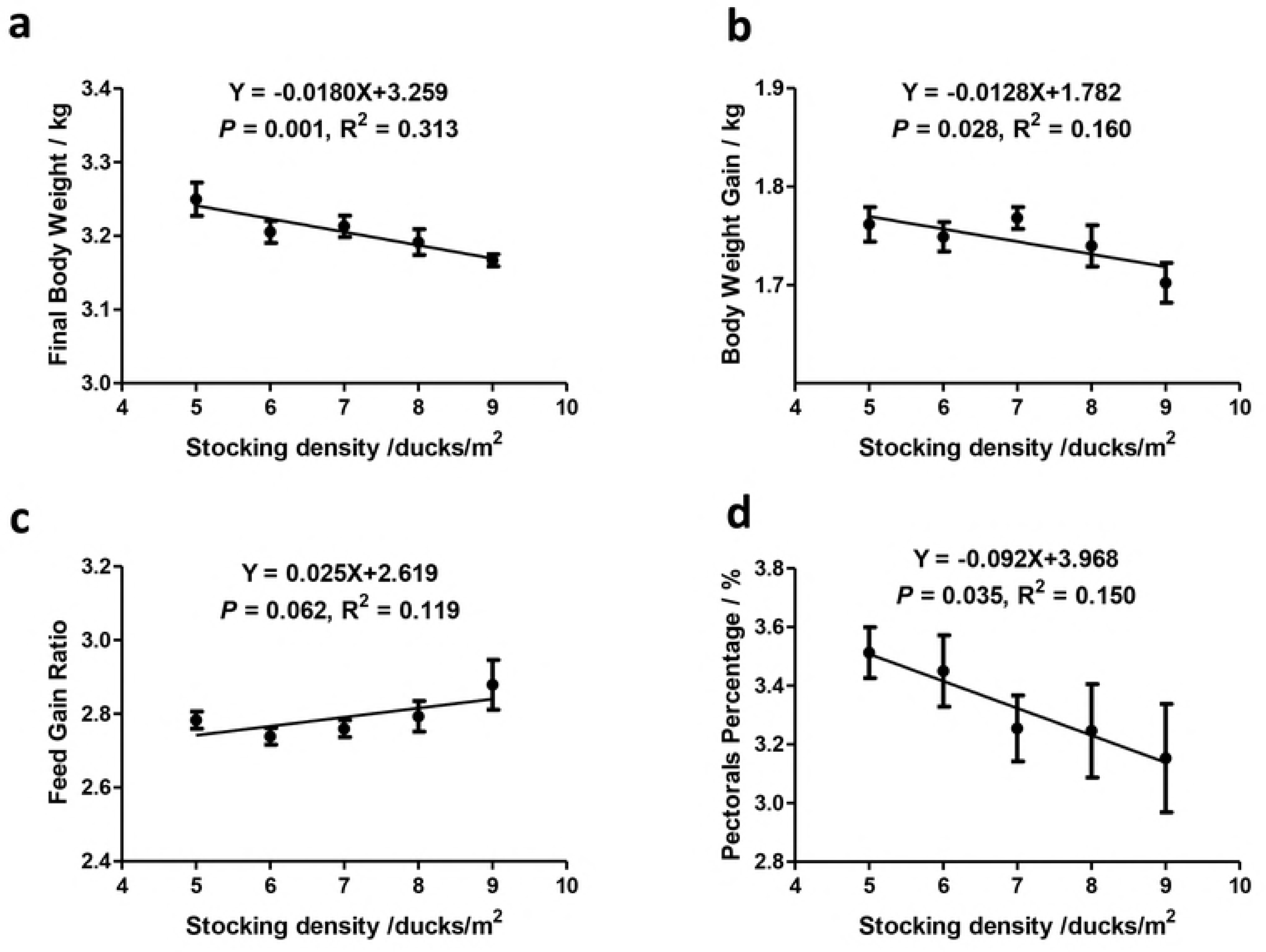
Duck performance and carcass traits. Data presented as means ± SE. n=6.

### Biochemical indices of serum and pectorals

Both redox and lipid metabolism indices from serum and pectoral samples were affected by stocking density.

In redox indices, raising density was significantly associated serum MDA and LDH, as well as pectoral T-AOC (Table 1). Serum MDA increased linearly, while pectoral T-AOC decreased linearly with increasing stocking density (Table 1).

**Table 1.** Redox related biochemical indices in serum and pectorals.

|  | Pectorals |  | Serum |  |  |
| --- | --- | --- | --- | --- | --- |
| density/m<sup>2</sup> | T-AOC | MDA | T-AOC | MDA | LDH |
| 5 | 0.71±0.15<sup>AB</sup> | 2.12±0.60 | 15.79±2.30 | 7.39±0.54<sup>B</sup> | 9186.19±888.64<sup>A</sup> |
| 6 | 0.50±0.14<sup>AB</sup> | 1.42±0.59 | 13.03±5.35 | 7.47±0.51<sup>B</sup> | 9224.32±195.84<sup>A</sup> |
| 7 | 0.48±0.28<sup>AB</sup> | 1.74±0.77 | 10.77±5.51 | 7.10±0.58<sup>B</sup> | 10192.19±732.17<sup>A</sup> |
| 8 | 0.46±0.16<sup>AB</sup> | 1.37±0.44 | 14.55±1.23 | 8.68±0.05<sup>AB</sup> | 7864.87±814.26<sup>B</sup> |
| 9 | 0.22±0.08<sup>B</sup> | 1.87±0.46 | 11.51±3.95 | 9.57±1.56<sup>A</sup> | 9225.23±273.26<sup>A</sup> |
| *P*-value |  |  |  |  |  |
| ANOVA | 0.015 | 0.176 | 0.278 | 0.008 | 0.001 |
| Linear | 0.023 | 0.490 | 0.253 | 0.038 | 0.238 |
| Quadratic | 0.069 | 0.148 | 0.269 | 0.068 | 0.470 |

Note: Total antioxidant capacity (T-AOC), malondialdehyde (MDA), lactate dehydrogenase (LDH).

Stocking density was also significantly altered lipid metabolism indices including serum LPL, TG, TC, HDL-C and LDL-C, as well as pectoral LPL (Table 2). Additionally, serum LDL-C increased linearly, and HDL-C had a linear increasing trend with increasing density. Serum TG and pectoral LPL had quadratic trends associated with stocking density (Table 2).

**Table 2.** Lipid metabolism related biochemical indices.

|  | Pectorals | Serum |  |  |  |  |
| --- | --- | --- | --- | --- | --- | --- |
| Group | LPL | LPL | TG | TC | HDL-C | LDL-C |
| 5 | 0.67±0.10<sup>BC</sup> | 2.76±0.62<sup>A</sup> | 1.01±0.22<sup>C</sup> | 4.14±0.32<sup>B</sup> | 4.01±0.30 | 3.24±0.60<sup>B</sup> |
| 6 | 0.99±0.23<sup>AB</sup> | 1.15±0.17<sup>B</sup> | 1.54±0.07<sup>B</sup> | 4.17±0.30<sup>B</sup> | 4.76±0.68 | 3.59±0.59<sup>B</sup> |
| 7 | 1.13±0.27<sup>A</sup> | 2.67±0.39<sup>A</sup> | 1.31±0.12<sup>BC</sup> | 4.13±0.15<sup>B</sup> | 4.41±0.83 | 4.83±0.73<sup>A</sup> |
| 8 | 0.84±0.30<sup>ABC</sup> | 2.52±0.79<sup>AB</sup> | 2.09±0.20<sup>A</sup> | 5.03±0.36<sup>A</sup> | 5.10±0.84 | 4.88±0.85<sup>A</sup> |
| 9 | 0.55±0.09<sup>C</sup> | 2.80±0.36<sup>A</sup> | 1.58±0.16<sup>B</sup> | 4.74±0.10<sup>A</sup> | 5.30±0.11 | 5.02±0.69<sup>A</sup> |
| *P*-value |  |  |  |  |  |  |
| ANOVA | 0.003 | <0.001 | <0.001 | <0.001 | 0.111 | 0.001 |
| Linear | 0.990 | 0.311 | 0.089 | 0.118 | 0.095 | <0.001 |
| Quadratic | 0.098 | 0.506 | 0.050 | 0.302 | 0.250 | <0.001 |

Note: lipoprotein lipase (LPL) triglyceride (TG), total cholesterol (TC), high-density lipoprotein cholesterol (HDL-C) and low-density lipoprotein cholesterol (LDL-C).

### Proteins identification and comparison

In GO analysis, the most striking different pathways between H and L group were small molecule metabolic process, organic acid metabolic process, oxoacid metabolic process and carboxylic acid metabolic process. Pathways including oxidation-reduction process, cellular amino acid metabolic process and fatty acid metabolic process were significant altered as well (Fig 2). Many proteins involved in redox, lipid metabolism, protein turnover, DNA repair, and immunity were associated with stocking density (Table 3).

**Fig 2.**
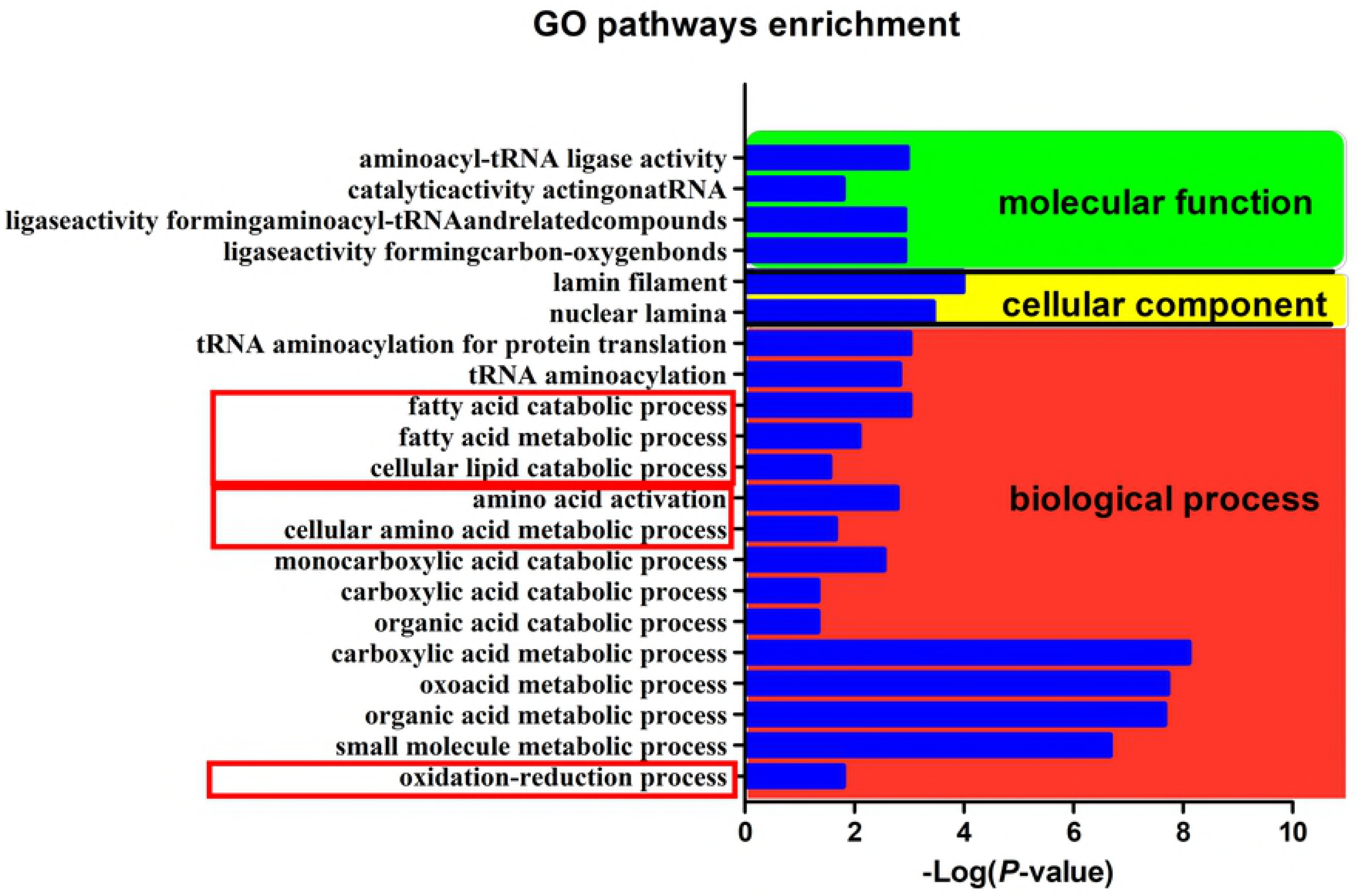
GO pathways enriched by H group. Log-transformed *P*-value is indicated the degree of difference between H group and L group.

**Table 3.** Differentially expressed proteins identified under different stocking density.

| Accession ID<sup>a</sup> | Annotation | Gene | H:L<sup>b</sup> | P-value |
| --- | --- | --- | --- | --- |
|  | **Redox related proteins** |  |  |  |
| U3I9D6 | DDRGK domain containing 1 | DDRGK | 1.218 | 0.010 |
| MT | Metallothionein | N/A | 1.332 | 0.014 |
| U3ID81 | Regucalcin | RGN | 0.706 | 0.025 |
| R0M3U9 | Catalase | Anapl_09675 | 0.882 | 0.051 |
|  | **Lipid metabolism related proteins** |  |  |  |
| U3IAY7 | Acyl-CoA dehydrogenase, long chain | ACADL | 1.154 | 0.040 |
| U3I7T9 | Acyl-CoA synthetase family member 2 | ACSF2 | 1.103 | 0.032 |
| U3J928 | Acyl-coenzyme A oxidase | ACOX1 | 1.098 | 0.007 |
| U3I7I1 | 2-hydroxyacyl-CoA lyase 1 | HACL1 | 1.143 | 0.095 |
|  | **Protein turnover related proteins** |  |  |  |
| U3I9U9 | Pyridoxal kinase | PDXK | 1.324 | 0.011 |
| U3IHG9 | Ribosomal protein S20 | RPS20 | 1.103 | 0.045 |
| U3I350 | Aspartyl-tRNA synthetase | DARS | 1.119 | 0.096 |
| U3J7Z7 | Asparaginyl-tRNA synthetase | NARS | 0.904 | 0.068 |
|  | **DNA repair related proteins** |  |  |  |
| U3IUN7 | RNA binding motif protein, X-linked | RBMX | 1.136 | 0.068 |
| U3IUB7 | Lamin B2 | LMNB2 | 1.119 | 0.076 |
|  | **Immunity related proteins** |  |  |  |
| R0K747 | Macrophage mannose receptor 1 | Anapl_08603 | 1.170 | 0.004 |
| Q6JWQ2 | MHC class I antigen alpha chain | Anpl-U | 0.785 | 0.021 |

Antioxidant related proteins such as regucalcin and catalase were downregulated in H group. Oxidative response related proteins like DDRGK domain containing 1 and metallothionein were enhanced in H group.

Fatty acid synthesis protein acyl-CoA synthetase family member 2 (ACSF2) was enriched in high stockind density. Fatty acid oxidation connected proteins including acyl-CoA dehydrogenase long chain (ACADL), acyl-coenzyme A oxidase (ACOX1) and 2-hydroxyacyl-CoA lyase 1 (HACL1) were enriched in the H group.

Protein turnover related proteins including Ribosomal protein S20, Pyridoxal kinase and aspartyl-tRNA synthetase (DARS2) were elevated while Asparaginyl-tRNA synthetase was reduced in H group.

In DNA repair involved proteins, Both RNA-binding motif protein X-linked (RBMX) and Lamin B2 showed enhancing trends in H group. The immunity associated macrophage mannose receptor (MR) 1 was elevated, while Major histocompatibility complex (MHC) class I antigen α chain was decreased in the H group.

### 16S rDNA analysis of cecal microbiota

Microbiota PCA analysis showed a clear separation of samples from the H group and samples from the L group (Fig 3a). The H group and L group had 290 bacterial taxa in common, while the H group and L group had 35 and 13 unique taxa, respectively (Fig 3b).

**Fig. 3.**
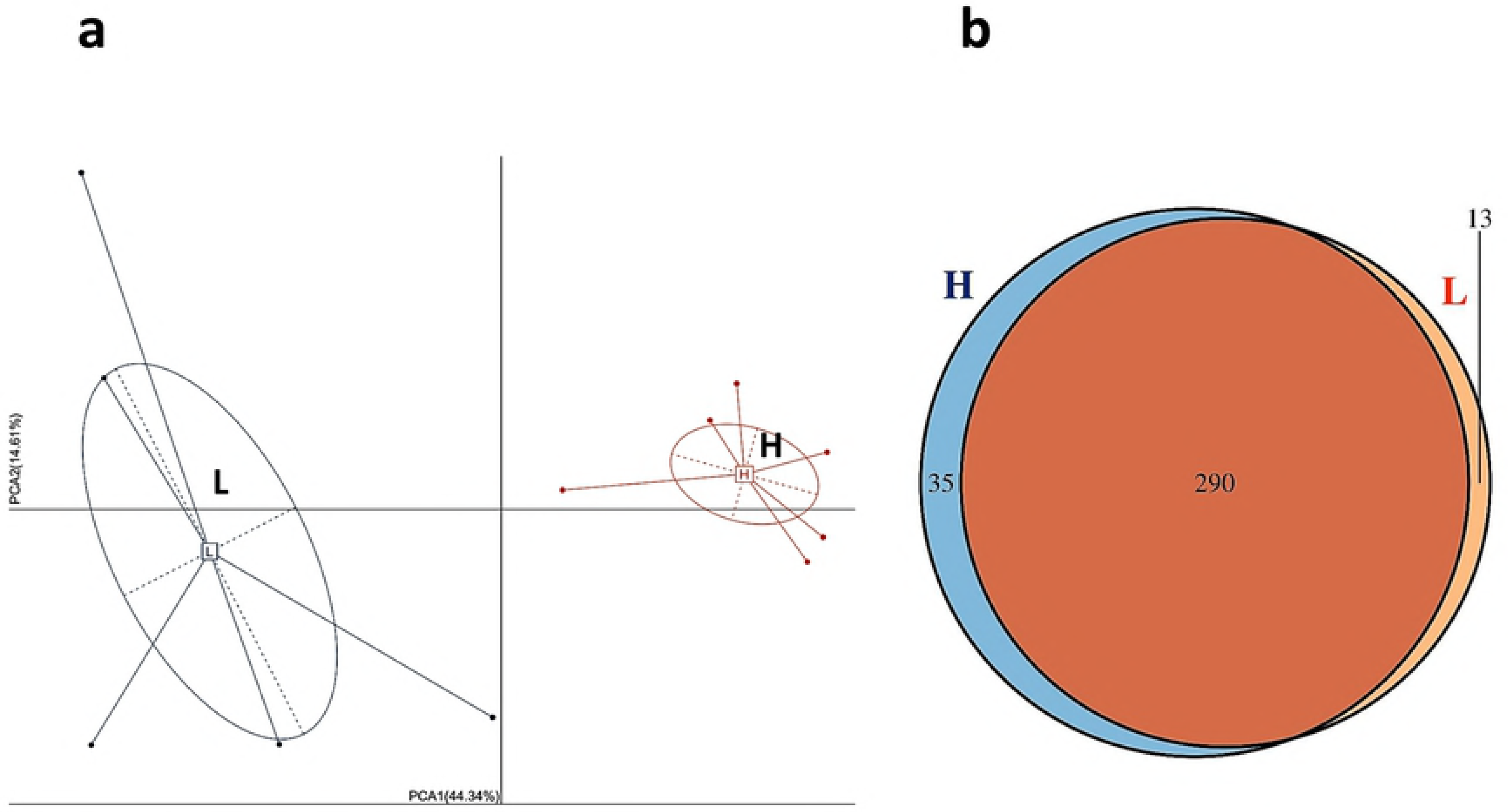
PCA and Veen chart of microbiota. a, PCA diagram; b, Veen chart.

LEfSe analysis, revealed higher relative abundance of *Firmicutes* and *Phascolarctobacterium* in the H group, while *Bacteroidia, Bacteroidales, Thermoplasmata, Methanomassiliicoccales, Methanomassiliicoccus* and *Methanomassiliicoccaceae* were more abundant in the L group (Fig 4a, 4b). Additionally, the H group had elevated *Lachnospiraceae* and *Ruminococcaceae* lower abundance of *Butyricimonas, Desulfovibrionaceae, Alistipes* and *Clostridiales* (Fig 5). Overall, the ratio of *Firmicutes* to *Bacteroidetes* in the H group was higher than in the L group (*P* = 0.058).

**Fig 4.**
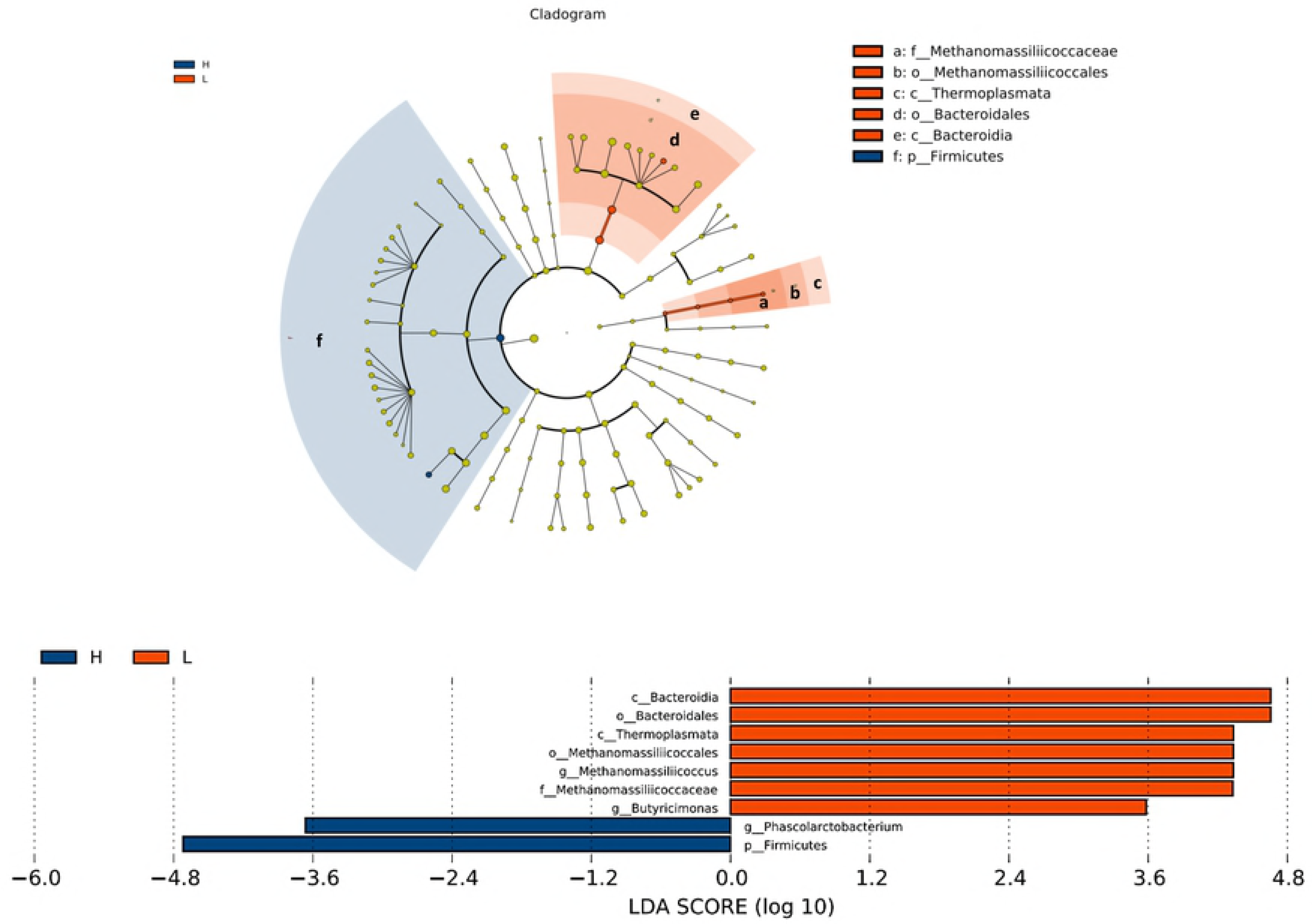
Bacterial taxa in H or L groups by LEfSe analysis. Phylogenetic relationships among significant bacterial biomarkers are indicated in the cladogram (top). Log-transformed linear discriminant analysis (LDA) scores of the significant biomarkers are indicated the bar chart (bottom).

**Fig 5.**
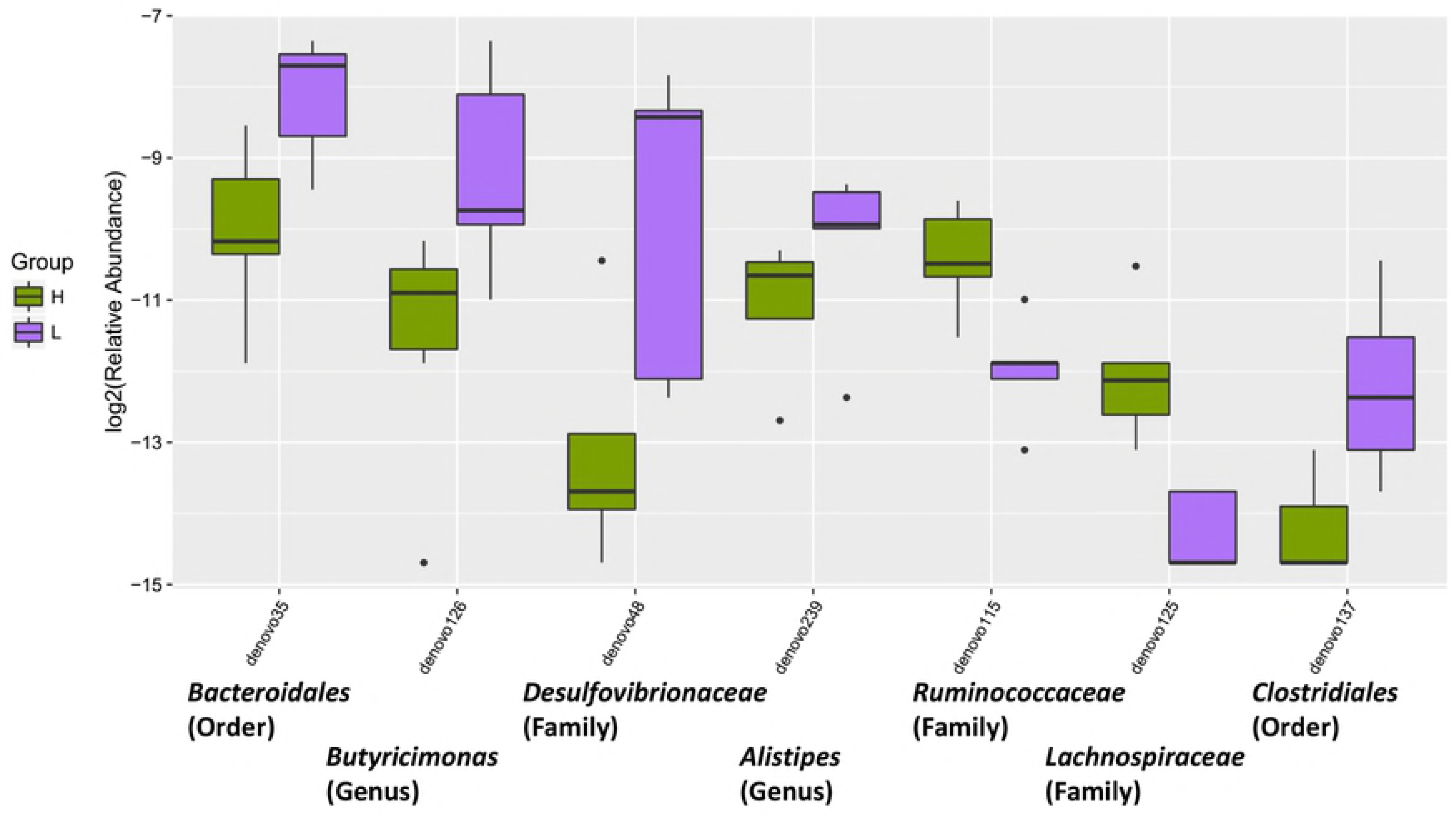
Relative abundance of anti-inflammatory bacterial taxa in H or L groups.

## DISCUSSION

Previous studies have indicated that that high raising density has negative influences on physiology and overall meat production [2-4]. This study showed a linear decrease of BW with increasing stocking density. Similarly, breast muscle was depressed by increasing stocking density. Which was consistent with the formerly research that increasing stocking density decreased breast fillet weight and its relative yield [18].

Increased density is associated with impaired antioxidant capacity, including decreasing total glutathione concentration and the glutathione (GSH): oxidized glutathione (GSSG) ratio [2]. Liver antioxidants including superoxide dismutase (SOD), catalase (CAT), GSH have previously been found to be reduced under high raising density [19]. In this study, enhancing stocking density increased serum MDA and decreased pectoral T-AOC. In proteomic analysis of this study, regucalcin and catalase expression was reduced in high density group. Catalases are involved in protecting cells from the damaging effects of ROS [20], while regucalcin is a calcium-binding protein with multiple roles that include calcium homeostasis, anti-oxidative, anti-apoptosis, and anti-proliferation [21]. Decreases in these proteins indicates that high stocking density may reduce duck anti-oxidant capacity.

High stocking density increases blood oxidative stress, including acute phase proteins, heat shock protein 70, and circulating corticosterone [22]. In current study, high stocking density had higher expressions of DDRGK domain containing 1, metallothionein, DARS2 and RBMX. DDRGK domain containing 1 and metallothionein were enriched in high density group. DDRGK1 is an endoplasmic reticulum membrane protein and plays an important role in maintaining the homeostasis of endoplasmic reticulum [23]. Metallothionein is an essential protein for the protection of cells against reactive oxygen species (ROS) [24]. Therefore, metallothionein can neutralize ROS and protect host from oxidative stress. A previous study found that loss of mitochondrial DARS2 leads to the activation of various stress responses in a tissue-specific manner independently of respiratory chain deficiency [25]. The RBMX is a nuclear protein that is involved in alternative splicing of RNA [26]. It is able to confer resistance to DNA damage [27]. Enhancement of these proteins suggests oxidative stress of Peking ducks under high stocking density.

Lipid storage is essential for protection against ROS toxicity [28]. The lipogenesis occurs in liver most exclusively, especially to waterfowls [29]. High stocking density was previously associated higher hepatic TG storage [30]. This study showed serum LDL-C increased with density incrementing. Moreover, high density group elevated ACSF2 expression. Acetyl-CoA synthetase catalyzes the formation of acetyl-CoA from acetate, CoA and ATP and participates in various metabolic pathways, including fatty acid and cholesterol synthesis and the tricarboxylic acid cycle [31]. ACSF2 belongs to the acyl-CoA synthetase family, activating fatty acids by forming a thioester bond with CoA [32]. Therefore, the increasing lipid biosynthesis may be the self-protection mechanism of Peking ducks under oxidative stress.

ROS are considered to be involved in the progression of non-alcoholic fatty liver disease (NAFLD) [33]. Interestingly, fatty liver production in waterfowls are fairly similar to human NAFLD [34]. Therefore, waterfowls can be a good model for liver steatosis research. In current study, ACADL, ACOX1 and HACL1 were enriched in high stocking density group. ACADL is a key enzyme participating in fatty acid oxidation [35]. Similary, ACOX1 are involved in β-oxidation in the liver [36]. HACL1 has two important roles in α-oxidation, the degradation of phytanic acid and shortening of 2-hydroxy long-chain fatty acids so that they can be degraded further by β-oxidation [37]. It has been reported that inhibition of β-oxidation decreases NADPH levels and increases ROS levels [38]. Therefore, elevation of these proteins may protect ducks from oxidative stress. It is increasingly recognized that the composition of the gut microbiota plays a critical role in influencing predisposition to chronic liver disorders such as NAFLD [12]. Samples from high stocking density had high levels of *Lachnospiraceae.* Interestingly, high *Lachnospiraceae* abundance was observed in patients with nonalcoholic steatohepatitis [39]. *Ruminococcaceae* and *Alistipes* were depleted in the high density group. Cirrhotic patients were previously shown to have lower *Ruminococcaceae* (7α-dehydroxylating bacteria) abundance compared to healthy patients [40]. In addition, *Alistipes* was significantly more abundant in the gut microbiota of healthy subjects compared to NAFLD patients [41].

Immunity can be regulated by oxidative stress. A decreased broiler bursa weight was previously reported to be associated with higher stocking densities [2]. DDRGK domain-containing protein 1 and Macrophage mannose receptor (MR) expression was higher in high density compared to the low density group. DDRGK1 is also an important regulatory protein of NF-κB [42]. Recent study found MR can protect against ROS burst [43]. Increase of these proteins reflects the immune response status Peking ducks under high stocking density. MHC class I antigen α chain was decreased under high stocking density. MHC class I plays a crucial role in immunity by capturing peptides for presentation to T cells and natural killer (NK) cells [44]. Dysregulation of MHC class I was correlated with unfolded protein response (UPR) and endoplasmic reticulum (ER) stress [45]. Upregulation of these proteins in the high density group suggests the immune adaption to high stocking. Moreover, high stocking density has been previously associated with adverse effects on the chicken intestinal commensal bacteria [4]. In the current study, *Phascolarctobacterium* was enriched, while *Bacteroidales, Butyricimonas* and *Alistipe* are depleted in high density group. A previous study found that *Phascolarctobacterium* was significantly correlated with systemic inflammatory cytokines [46]. Besides, depletion of *Bacteroidales* has previously been associated with disease status [47]. Additionally, reduction in *Butyricimonas* is associated with increased proinflammatory gene expression [48]. Studies confirm that patients with IBD and *Clostridium difcile* infection have a lower abundance of *Alistipe* than their healthy counterparts [49]. The current study showed higher *Firmicutes* to *Bacteroidetes* ratio in high density group. The microbiota of irritable bowel syndrome (IBS) patients, compared with controls, had a 2-fold increased ratio of the *Firmicutes* to *Bacteroidetes* [50]. Bile acids (BAs) are important metabolites of the microbiome and can modulate the composition of the gut microbiota directly or indirectly through activation of the innate immune system [51]. Bile acid plays a crucial role in control of inflammation and NAFLD [52]. Samples from the high stocking density group had higher levels of *Ruminococcaceae* and lower abundance of *Desulfovibrionaceae*, and *Clostridiales* compared to low density group. Cholesterol 7α-hydroxylase (CYP7A1) is the enzyme responsible for catalyzing the first and rate-limiting step in the classical bile acid synthetic pathway [53]. Futhermore, an inverse relationship between Cyp7a1 expression and *Ruminococcaceae* abundance has been previously demonstrated [19]. A study found enriched *Desulfovibrionaceae* was accompanied by increased hepatic taurine-conjugated cholic acid and β-muricholic acid, which were the main constituent of bile acid pool [54]. *Clostridiale* was positively correlated with muricholic acid as well [19]. A loss of bacteria belonging to the *Clostridiales* order was correlated with a disturbance in the bile-microbial axis [55]. Therefore, alternations of bacteria showed a decrease trend in bile acid synthesis.

In conclusion, high stocking density caused oxidative stress, which involved in alternation of gut-liver axis (Fig 6).

**Fig 6.**
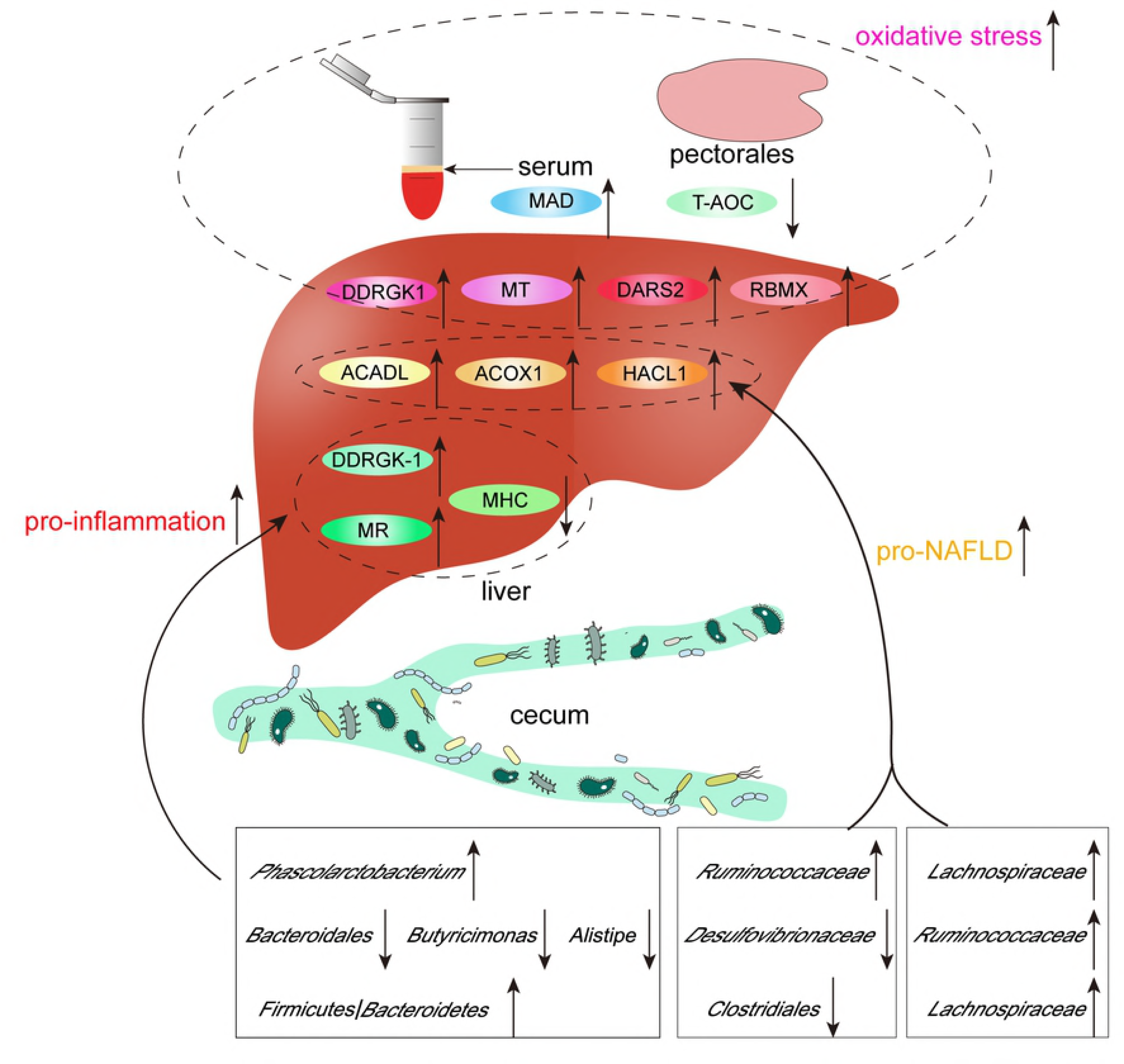
Graphic summary of liver proteome and gut microbiota alternations under high stocking density.

## Acknowledgements

This research was supported by the System for Poultry Production Technology, Beijing Agriculture Innovation Consortium (Project Number: BAIC04-2017).

